# Clade-defining mutations in human H1N1 hemagglutinin protein from 2021-2023 have opposing effects on *in vitro* fitness and antigenic drift

**DOI:** 10.1101/2024.05.18.594815

**Authors:** Nicholas J Swanson, Jyothsna Girish, Madeline Yunker, Hsuan Liu, Julie Norton, Christa Han, Heba Mostafa, Katherine Fenstermacher, Richard Rothman, Andrew Pekosz

**Author notes:** Corresponding Author: Andrew Pekosz.

## Abstract

Seasonal influenza viruses frequently acquire mutations that have the potential to alter both virus replication and antigenic profile. Recent seasonal H1N1 viruses have acquired mutations to their hemagglutinin (HA) protein receptor binding site (RBS) and antigenic sites, and have branched into the clades 5a.2a and 5a.2a.1. Both clades demonstrated improved *in vitro* fitness compared with the parental 5a.2 clade as measured through plaque formation, infectious virus production in human nasal epithelial cells, and receptor binding diversity. Both clades also showed reduced neutralization by serum from healthcare workers vaccinated in the 2022-23 Northern Hemisphere influenza season compared to the vaccine strain. To investigate the phenotypic impact of individual clade-defining mutations, recombinant viruses containing single HA mutations were generated on a 5a.2 genetic background. The 5a.2a mutation Q189E improved plaque formation and virus replication, but was more efficiently neutralized by serum from individuals vaccinated in 2022-23. In contrast, the 5a.2a mutation E224A and both 5a.2a.1 mutations P137S and K142R impaired aspects of *in vitro* fitness but contributed significantly to antigenic drift. Surprisingly, the E224A mutation and not Q189E caused broader receptor binding diversity seen in clinical isolates of 5a.2a and 5a.2a.1, suggesting that receptor binding diversity alone may not be responsible for the phenotypic effects of the Q189E mutation. These data document an evolutionary trade-off between mutations that improve viral fitness and those that allow for the evasion of existing host immunity.

**Importance:** Seasonal influenza is a significant contributor to global morbidity and mortality. To better understand how these viruses evolve, recent seasonal H1N1 viruses were characterized and the impact of individual mutational differences between the hemagglutinin proteins of these viruses was assessed. Mutations that contribute to escape from preexisting immunity were deleterious to virus replication in human nasal epithelial cell cultures, requiring the accumulation of additional mutations to compensate for this loss of fitness. This adds to our understanding of how influenza evolution balances replication fitness with antigenic drift.

## Introduction

Seasonal influenza viruses are among the largest causes of global respiratory infections. Interventions including antivirals and vaccinations have helped reduce the burden of disease, but influenza continues to infect roughly one billion people and cause the deaths of hundreds of thousands each year (1). One of the major challenges to combatting this disease is the genetic diversity present within these viruses, and the rapid pace of continued diversification. “Seasonal influenza” refers to two co-circulating genera of viruses from the *Orthomyxoviridae* family, influenza A virus (IAV) and influenza B virus, which can each be further subdivided into strains and subtypes. IAV is the predominant contributor to annual disease burden (2), and within IAVs both H1N1 and H3N2 subtypes co-circulate with relative prevalence changing from season to season (3).

Each IAV subtype consists of multiple clades as defined by the hemagglutinin (HA) sequence, with distinct genotypes and antigenic profiles (4). Mutations can alter numerous aspects of viral phenotype, including changes to both fitness and antigenic profile. Changes to fitness can include differences in receptor binding (5–8), virus replication (8–10), interactions with innate immunity (11), transmissibility (12–14), and other characteristics (15). Antigenic profile refers to the regions of a virus which are recognized by host antibodies, and changes to this profile through antigenic drift can alter how the virus appears to the immune system and result in a loss of antibody binding. Frequent mutations to viral antigenic sites contribute to our need for annual vaccination with updated strains whose antigenic profiles match those of currently circulating viruses (16). HA functions as the primary antigenic determinant of influenza and is also responsible for receptor binding and virus-host membrane fusion (17). These roles result in dual selective pressures on influenza to both evade existing host immunity and maintain or improve virological fitness by optimizing cell attachment and entry.

The H1N1 subtype has seen significant genetic diversification over the past several seasons, with multiple clades emerging and achieving varying degrees of prevalence. In the 2019-20 Northern Hemisphere (NH) influenza season the 5a.1 and 5a.2 clades both emerged from the 5a parental clade (18). The 5a.1 clade quickly predominated over 5a.2, and the 5a.2 clade was shown to have impaired *in vitro* fitness compared with 5a.1 which may have contributed to its limited spread (8).The 5a.2 clade rapidly acquired a series of HA mutations at the start of 2021 and branched into a novel subclade, 5a.2a, which quickly became the predominant H1N1 clade. 5a.2a is defined by mutations in the receptor binding site and antigenic site mutations A186T, Q189E, and E224A (H1 numbering). The A186T mutation has emerged previously in regionally circulating H1N1, including Ukraine (19), Tunisia (20), Turkey (21), Spain (22), Cuba (23), and the Czech Republic (24). Notably, Q189E had appeared as a clade-defining mutation in 5a.1 (8) but is new to the 5a.2 lineage. The E224A mutation has been shown to affect viral pathogenesis (25), cell tropism (26), and receptor binding (27, 28), and alanine already exists as the predominant residue at site 224 in swine H1 HA (25). Initial serological characterization indicates that 5a.2a is antigenically drifted from the 5a.2 clade (29, 30).

As the 5a.2a clade circulated in 2021 it continued to acquire mutations, and by 2022 the subclade 5a.2a.1 had emerged but remained a minority clade. It is defined by mutations including P137S and K142R, which occupy both the receptor binding site and antigenic site Ca2. Both of these mutations have previously emerged in regional circulation (22, 31–33), and have been identified repeatedly in circulating swine H1 HA (34–37). Serological data suggests that this clade may be antigenically drifted from both the 2022-23 NH influenza vaccine strain and the 5a.2a clade (30).

To evaluate phenotypic changes to these recent H1N1 clades, experiments were performed to identify both antigenic and *in vitro* fitness differences between representative virus genotypes. Both 5a.2a and 5a.2a.1 clades were antigenically drifted from 5a.2 and demonstrated improved *in vitro* fitness. The mutations contributing to escape from vaccine induced immunity did so at the expense of infectious virus production and a mutation that improved virus replication increased virus recognition by serum from vaccinated individuals. These data highlight the evolutionary trade-off between acquiring mutations that allow for the evasion of existing immunity, while maintaining adequate virological fitness.

## Results

### Emergence of novel 5a.2 subclades

After the emergence of the 5a.1 and 5a.2 clades in 2019, 5a.1 quickly predominated and almost completely overtook 5a.2 by the end of 2020 (Figure 1A). Characterization of these viruses revealed a reduction in *in vitro* fitness for 5a.2 compared with 5a.1 (8), which may have contributed to the success of the latter. In 2021, a series of mutations emerged within the 5a.2 clade which were subsequently classified as a new clade, 5a.2a. 5a.2a spread rapidly and by 2022 overtook 5a.1 to become the primary H1N1 clade (Figure 1A). The mutations defining this new 5a.2a clade include multiple changes to antigenic sites, as well as several differences in the receptor binding site of the HA protein (Figure 1B). This suggests potential adaptations to the viral phenotype which may have improved fitness or antigenic profile, allowing it to circulate more widely than its 5a.2 parental clade.

**Figure 1:**
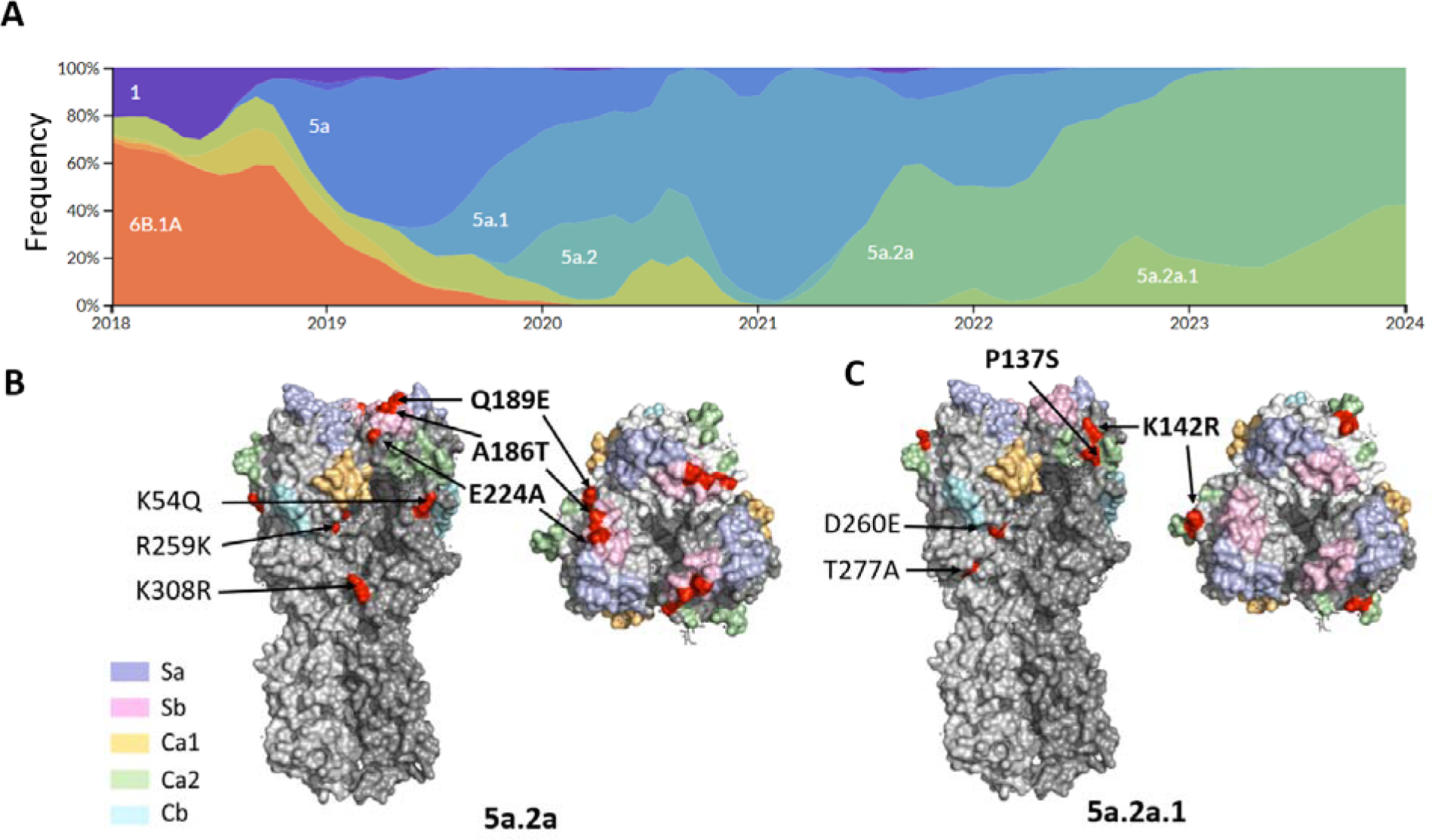
Emergence of seasonal H1N1 clades. A) Frequencies of circulating clades of H1N1 since 2018, generated using the NextStrain pipeline (47). Clade 5a.2a emerged in 2021 and quickly predominated, followed by 5a.2a.1 which slowly gained prevalence. Clade-defining mutations of B) 5a.2a and C) 5a.2a.1 mapped on the hemagglutinin trimer using PyMOL (PDB: 3LZG), with antigenic sites shaded. In addition to the mutations shown, 5a.2a.1 also contains all the clade-defining mutations of 5a.2a. Mutations surrounding the receptor binding site are in bold.

The 5a.2a.1 clade, an offshoot of 5a.2a, also appeared around this time and slowly rose in prevalence as a sizable minority within H1N1 (Figure 1A). In addition to the mutations that define the 5a.2a clade, 5a.2a.1 contains several changes to both the receptor binding site and antigenic sites (Figure 1C), and initial serological data indicate that this clade may be antigenically drifted from 5a.2a (30), though its limited initial success suggests that any phenotypic changes acquired by this clade did not produce overwhelming advantages over 5a.2a.

To identify phenotypic differences between the 5a.2a and 5a.2a.1 clades, representative viruses were isolated from clinical samples of H1N1 collected from the Johns Hopkins Hospital during the 2022-23 Northern Hemisphere (NH) influenza season. These include two 5a.2a viruses, A/Baltimore/JH-22363/2022 and A/Baltimore/JH-22377/2022, and one 5a.2a.1 virus, A/Baltimore/JH-22400/2022. Two 5a.2a viruses were chosen to reflect intra-clade genetic diversity; A/Baltimore/JH-22377/2022 contains the T216A hemagglutinin mutation which appeared with regularity in this clade in 2022 and which has increased in prevalence in H1N1 since the 2022-23 season. In addition, a 5a.2 virus from the 2019-20 NH season, A/Baltimore/R0675/2019, was included as a comparison group for the parental 5a.2 clade.

### Recent H1N1 clades show improved *in vitro* fitness compared with the parental 5a.2 clade

To compare *in vitro* fitness between these clades, plaque formation and viral replication were assessed. Both 5a.2a and 5a.2a.1 viruses produced significantly larger plaques on MDCK cells compared with the 5a.2 parental clade (Figure 2A, 2B). Additionally, the 5a.2a virus A/Baltimore/JH-22363/2022 produced significantly larger plaques than the 5a.2a.1 virus A/Baltimore/JH-22400/2022, though neither were significantly different from the 5a.2a virus A/Baltimore/JH-22377/2022 which contains the T216A mutation (Figure 2B). Low-MOI growth curves were then performed on human nasal epithelial cell (hNEC) cultures, which are primary differentiated cell cultures grown at an air-liquid interface, and are an *in vitro* model of the upper respiratory tract (38–42). Both 5a.2a viruses, but not 5a.2a.1, replicated to significantly higher titers at multiple timepoints compared with 5a.2 (Figure 2C). To compare total infectious virus production, the area under the curve was calculated for each growth curve, which confirmed that 5a.2a, but not 5a.2a.1, produced significantly more infectious virus compared with 5a.2 (Figure 2D). Differences in replication were observed primarily at timepoints after peak virus titer had been reached, so to investigate this phenotype, virus production was separated into pre-peak and post-peak sections. Total virus production pre-peak was not significantly different between any of the clades (Figure 2E), however post-peak virus production was significantly higher for both recent H1N1 clades compared with the 5a.2 clade (Figure 2F).

**Figure 2:**
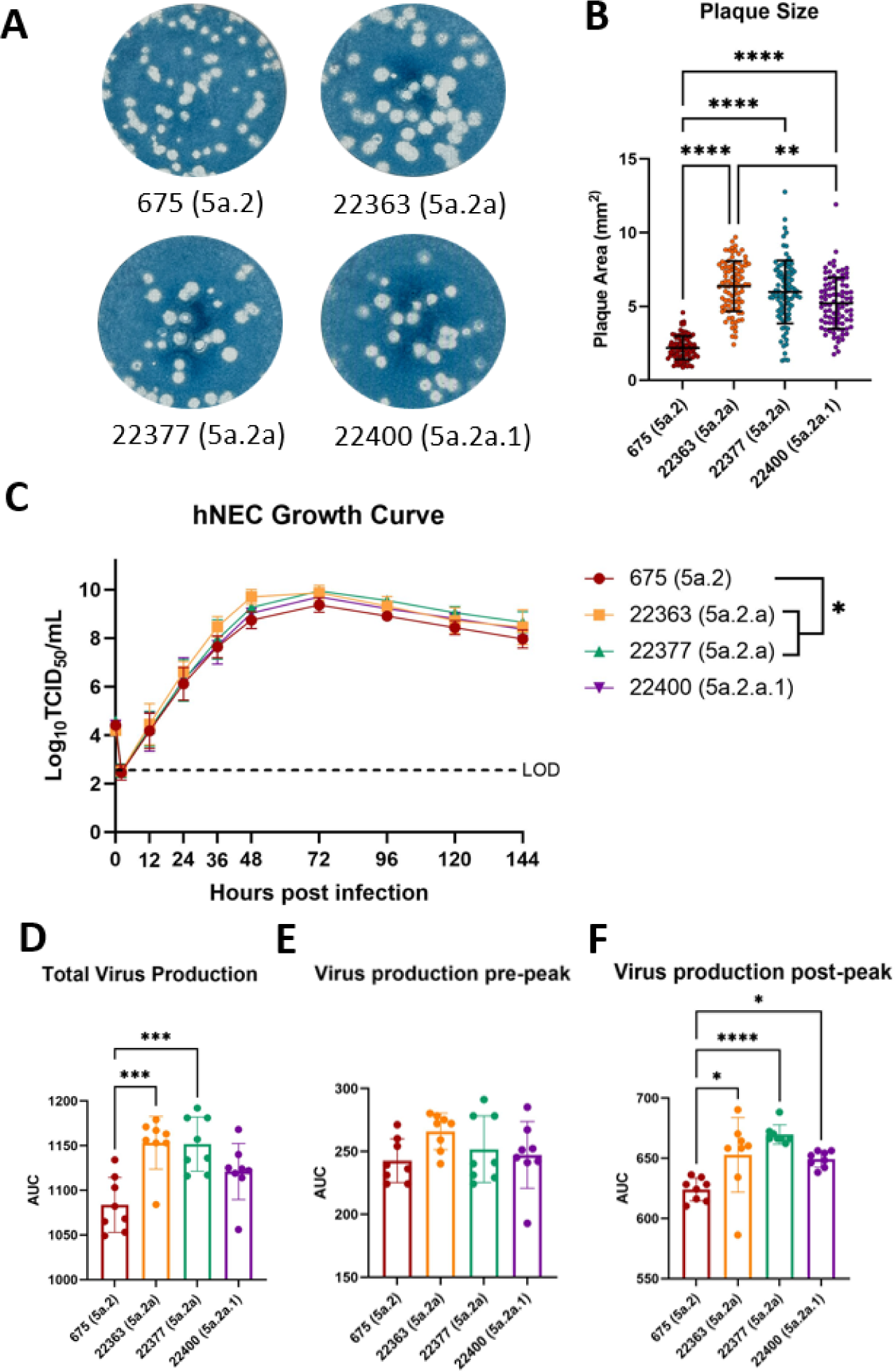
Characterization of *in vitro* replication fitness of circulating H1N1 viruses. A) Plaque formation on MDCK cells of representative viruses from circulating clades compared with their parental clade (5a.2). B) Recently circulating clades formed significantly larger plaques than their parental clade (Kruskal–Wallis ANOVA with Dunn’s multiple comparisons test). Growth curves on human nasal epithelial cells reveal an increase in C) the kinetics of infectious virus production and D) total virus production for the 5a.2a clade compared with 5a.2. Infectious virus production differences are not present in E) pre-peak titers (peak=72 hpi) but are driven by F) post-peak viral titers. One-way (AUC) or two-way (growth curve) ANOVA with Tukey post-hoc. * p<0.05, ** p<0.01, *** p<0.001, **** p<0.0001

### Recent H1N1 clades are antigenically drifted from their ancestral clade

Antigenic changes were assessed through virus neutralization assays using serum from the Johns Hopkins Centers for Excellence in Influenza Research and Response (JH-CEIRR) 2022 cohort. This cohort includes serum samples from healthcare workers both pre- and post-vaccination with the annual influenza vaccine, and participants include both men and women with ages ranging from 22-75. At both pre- and post-vaccination timepoints, the 5a.2a and 5a.2a.1 clades were less well neutralized by serum antibodies compared with the vaccine strain for that season, a virus from the 5a.2 clade (Figure 3A, 3B). The 5a.2a.1 clade also appeared to be antigenically drifted from 5a.2a, showing a significant reduction in neutralization both pre- and post-vaccination compared with at least one of the representative 5a.2a viruses. Interestingly, vaccination produced a comparable boost in neutralizing antibody titers across the virus panel; the fold change in neutralizing titers was similar for all four viruses tested (Figure 3C), and seroconversion rates were not significantly different (Figure 3D).

**Figure 3:**
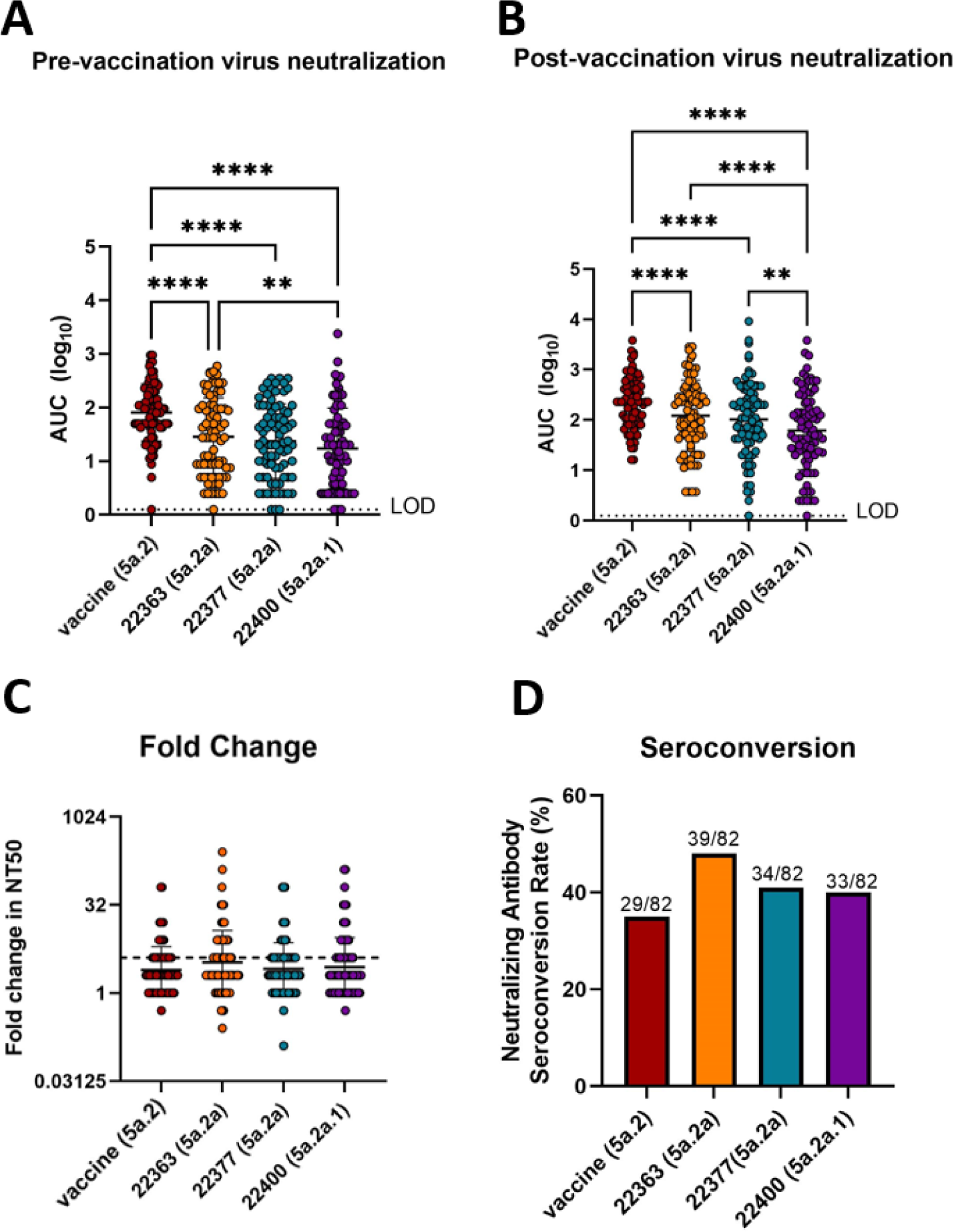
Neutralization of vaccine and circulating viruses by human serum. Virus neutralization was assessed using serum from a cohort of healthcare workers both A) pre- and B) post-vaccination with the 2022-23 Northern Hemisphere influenza vaccine. The virus panel included representative viruses from 5a.2a and 5a.2a.1 clades and the H1N1 vaccine strain, a 5a.2-clade virus (repeated measures one-way ANOVA with Tukey post-hoc). 5a.2a viruses were significantly less well neutralized than the vaccine virus, and 5a.2a.1 had a further reduction from 5a.2a in both serum panels. C) Fold change in neutralizing antibody titers after vaccination was similar for all viruses (repeated measures one-way ANOVA with Tukey post-hoc). D) Percent of serum donors who seroconverted (showed at least a fourfold increase in neutralizing titers post-vaccination) was similar for each virus (Chi-square test). N=82. ** p<0.01, **** p<0.0001

### Identifying the phenotypic contributions of individual clade-defining mutations

While the initial characterization of representative clinical isolates from these clades suggests important phenotypic changes associated with recent clade-defining mutations, the impact of individual gene segments and mutations cannot be determined by characterizing clinical virus isolates alone. To elucidate the contribution of individual genes and mutations, an infectious clone of the A/Baltimore/R0675/2019 5a.2 virus was generated as previously described (43) by creating eight plasmids containing individual gene segments of the virus (5, 44, 45). To confirm that the virus cloning and rescue process alone did not alter viral phenotype, a rescued A/Baltimore/R0675/2019 virus was compared with the genetically identical wildtype virus and both demonstrated similar *in vitro* fitness as judged by plaque sizes (Figure 4A and 4B) and replication on hNEC cultures (Figure 4C to 4F).

**Figure 4:**
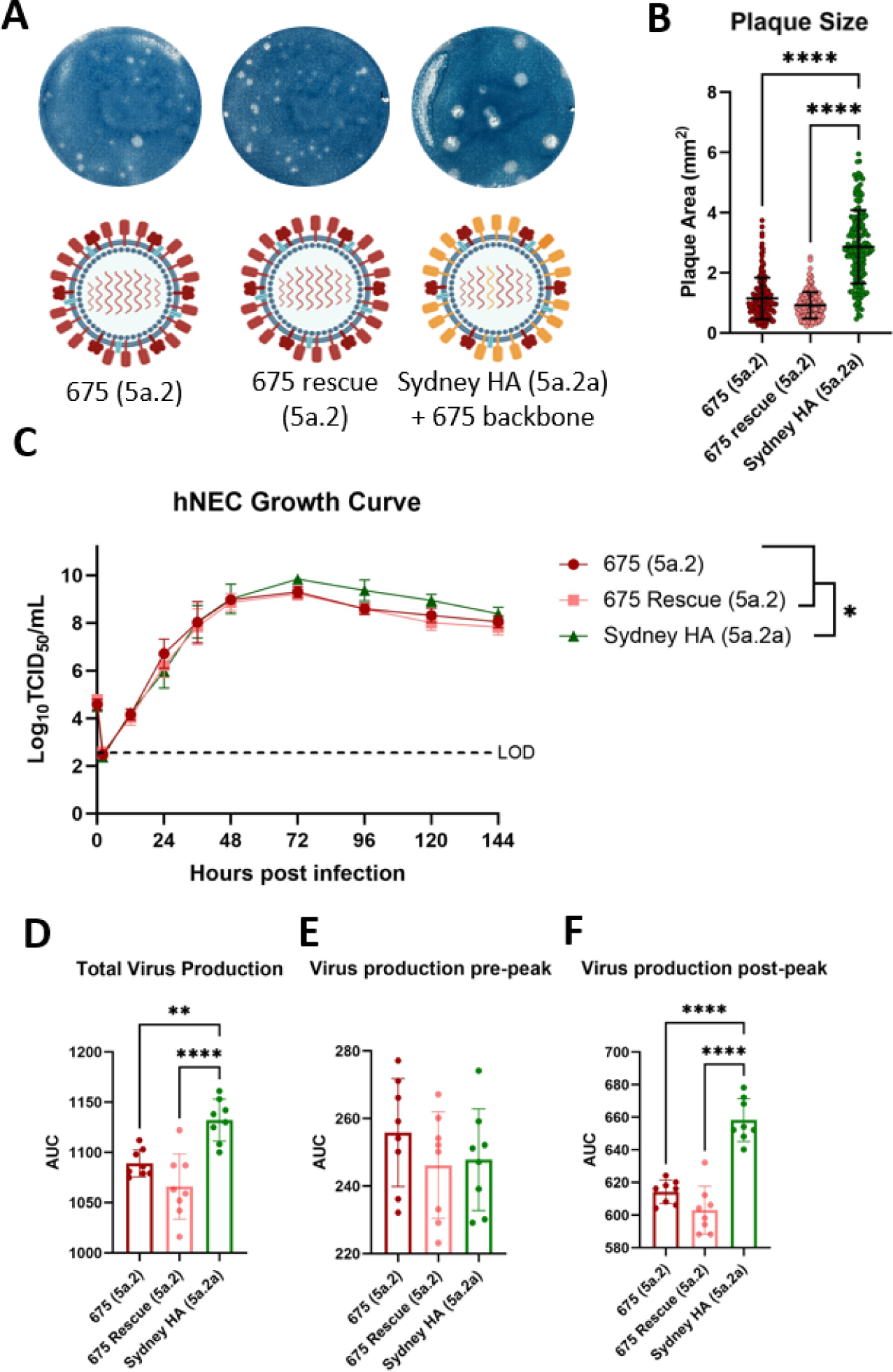
The contribution of the 5a.2a HA segment on *in vitro* replication fitness. A) A 1:7 reassortant virus was generated containing the HA from a 5a.2a virus (A/Sydney/5/2021) with all seven remaining segments from a 5a.2 virus (A/Baltimore/R0675/2019), and plaque assays were performed on MDCK cells. B) The reassortant virus produced significantly larger plaques than the wild type 5a.2 virus and a control rescue 5a.2 virus (Kruskal–Wallis ANOVA with Dunn’s multiple comparisons test). The reassortant virus replicated to a greater extent in hNEC cultures C), producing a greater amount of D) total infectious virus particles compared to the parental virus. Infectious virus production E) pre-peak was not different (peak=72 hpi) but more infectious virus was produced F) post-peak with the 5a.2a HA segment containing virus. One-way (AUC) or two-way (growth curve) ANOVA with Tukey post-hoc. ** p<0.01, **** p<0.0001

To confirm that changes to the hemagglutinin protein were responsible for the observed differences between 5a.2 and 5a.2a, a reassortant virus was generated containing seven gene segments from the 5a.2 virus and an HA segment from 5a.2a (A/Sydney/5/2021). This reassorted virus generated significantly larger plaques on MDCK cells (Figure 4A and 4B) and replicated to significantly higher titers on hNEC cultures compared with the 5a.2 virus (Figure 4C and 4D). Virus replication reflected the trend seen in clinical isolates, with differences in total virus production being driven by increases in post-peak virus production (Figure 4E and 4F).

After confirming that the HA protein was driving differences between clades, the contributions of individual clade-defining mutations were assessed. Viruses were generated using the seven segments from the 5a.2 virus along with 5a.2 HA segments containing individual point mutations from either the 5a.2a (A186T, Q189E, and E224A) or 5a.2a.1 (P137S and K142R) clades. The Q189E mutation from the 5a.2a clade significantly increased plaque size on MDCK cells compared with the 5a.2 virus, while the E224A mutation from 5a.2a and the P137S mutation from 5a.2a.1 both reduced plaque size (Figure 5A, 5B).

**Figure 5:**
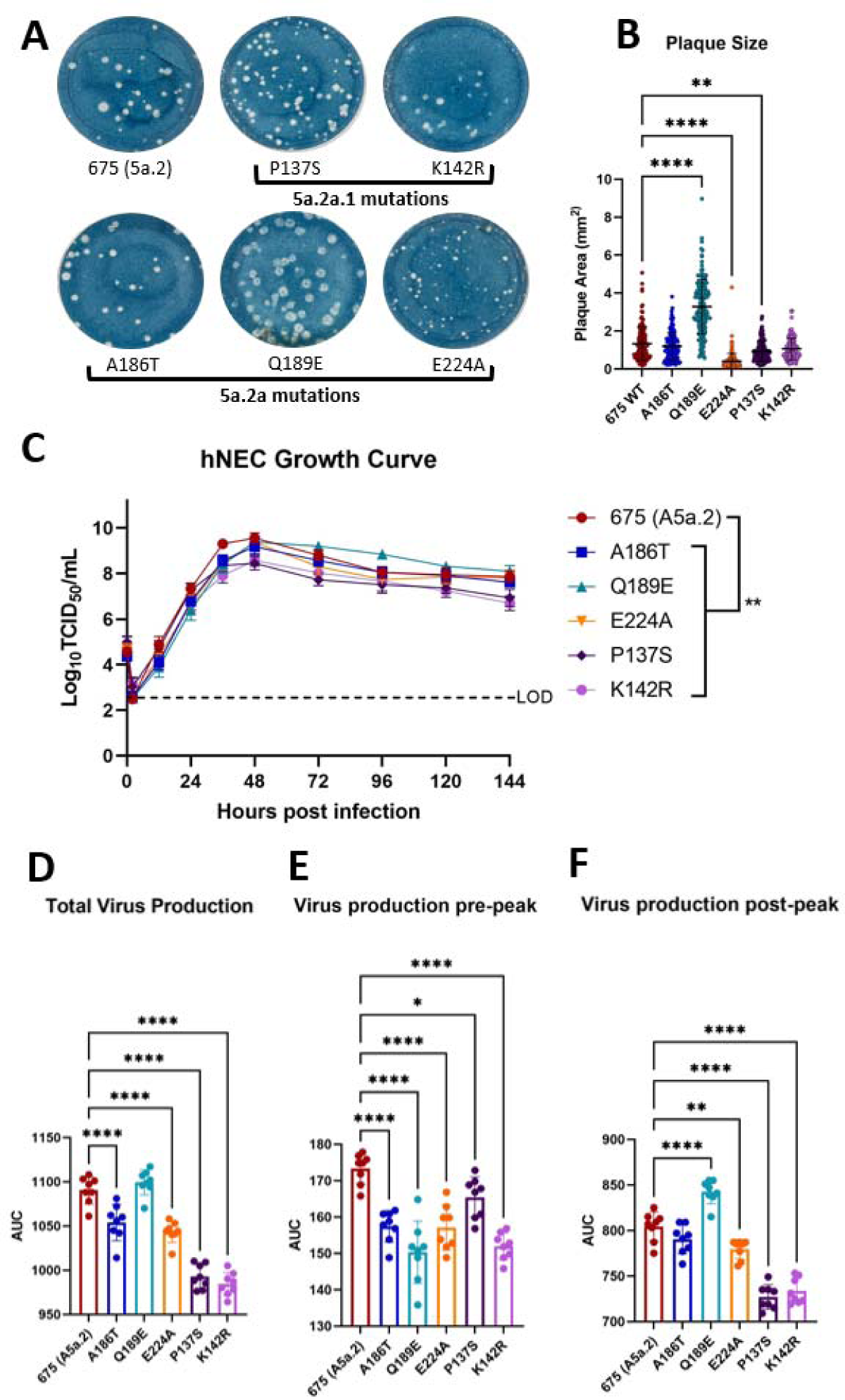
Determining the impact of individual clade-defining mutations in HA on *in vitro* replication fitness. Viruses were generated using a 5a.2 background (A/Baltimore/R0675/2019) containing individual point mutations from recent H1N1 clades (5a.2a and 5a.2a.1). A) Representative images of MDCK plaque assays. B) The Q189E mutation from 5a.2a significantly increased plaque area, while E224A and P137S reduced plaque area (Kruskal–Wallis ANOVA with Dunn’s multiple comparisons test). C) Viral growth curves on human nasal epithelial cells reveal significant temporal differences in replication between the 5a.2 virus and all point mutant viruses (Two-way ANOVA with Tukey post-hoc). D) Total virus production is reduced with all mutations except for Q189E. E) Pre-peak infectious virus titer is reduced for all point mutant viruses (peak=48 hpi), however virus production F) post-peak is only reduced for E224A, P137S, and K142R, while the Q189E mutation increases total virus production (One-way ANOVA with Tukey post-hoc). * p<0.05, ** p<0.01, *** p<0.001, **** p<0.0001

Growth curves on human nasal epithelial cells showed that every point mutant resulted in significantly different viral titers at multiple timepoints post-infection compared with the 5a.2 virus (Figure 5C). Area under the curve analysis revealed that total virus production was reduced for all viruses except for the Q189E mutant (Figure 5D), with every single mutant virus producing significantly less virus than the 5a.2 virus before peak titer (Figure 5E). Post-peak production, however, was increased for Q189E compared with 5a.2 (Figure 5F), mirroring the trend seen in both the clinical isolates and the 5a.2a reassorted virus. Both E224A and the two 5a.2a.1 mutations (P137S and K142R) produced significantly less virus post-peak compared with 5a.2 (Figure 5F).

### Differences in receptor binding between recent H1N1 clades

The changes observed to *in vitro* fitness of recent clades, combined with the location of clade-defining mutations on the receptor binding site, suggested a role for receptor binding as a mediator for fitness changes. To investigate receptor binding diversity, Consortium for Functional Glycomics glycan microarrays were performed on both the clinical isolate panel and the mutant virus panel. These glycan arrays contain over 100 terminally sialylated glycans including ones with either alpha 2,3 or alpha 2,6 linkages. Receptor binding profiles for both clinical viruses and point mutants demonstrate a preference for alpha 2,6 terminally sialylated glycans, the canonical human influenza receptor, and show differences in both quantity and identity of bound glycans (Figure 6A, 6B). To visualize these differences, a heat map was generated of all alpha 2,6 glycans included in the array, showing relative binding intensity for each virus. While there was some overlap in bound glycans among all viruses (indicated by a blue circle), both 5a.2a and 5a.2a.1 viruses bound to additional glycans not bound by the parental 5a.2 clade (indicated by a green circle), and these glycans were also bound by the E224A mutant virus (Figure 6C, CD). This suggests that the E224A mutation is responsible for this increase in receptor binding diversity of recent H1N1 clades. Additionally, the K142R mutation from the 5a.2a.1 clade was shown to abrogate binding to a specific glycan (ID number 560), which was strongly bound by all other viruses (indicated by a purple circle). The Q189E mutation that increased *in vitro* replication fitness did not show major changes in glycan binding pattern, suggesting that altered receptor binding diversity is most likely not the reason for the improved fitness of viruses containing this mutation. These results show that recent H1N1 mutations have produced altered receptor binding diversity but that receptor binding diversity alone does not correlate with overall viral fitness.

**Figure 6:**
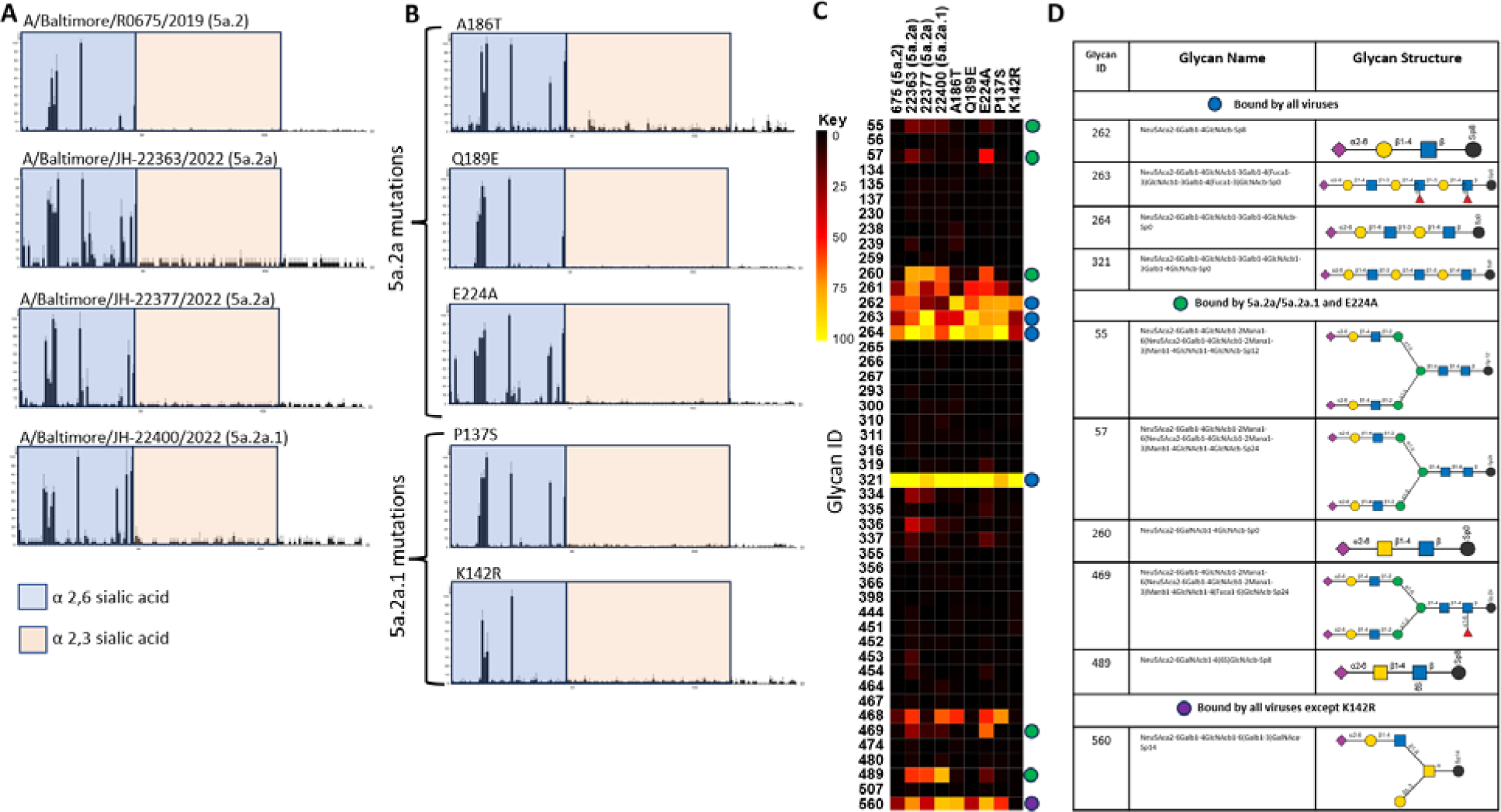
Comparison of receptor binding activity via the Consortium for Functional Glycomics (CFG version 5.5) glycan microarray. Receptor binding profiles of H1N1 clinical isolates A) and point mutants on a 5a.2-clade background B) show preference for alpha 2,6 terminally sialylated glycans, and differences in individual glycans bound. X axis depicts individual glycans included in the array, and Y axis depicts the relative fluorescence units (RFUs), a metric of virus binding, normalized to 100. C) heat map of normalized RFU values for each glycan, for each virus. A yellow square indicates maximum binding by that virus. Glycans of note are marked with colored circles; blue are glycans that were well-bound by all viruses, green are glycans bound well by viruses from the 5a.2a and 5a.2a.1 clades as well as the E224A point mutant virus, and purple is a glycan that was well-bound by all viruses except the K142R point mutant. D) Structures and names of glycans of note.

### Individual HA mutations can reduce or increase neutralization by serum from vaccinated individuals

To assess the contributions of individual clade-defining mutations on antigenic drift, neutralization assays were performed on the point mutant virus panel using post-vaccination serum from the 2022 JH-CEIRR healthcare worker cohort. The E224A, P137S, and K142R mutations all significantly reduced serum neutralization compared with the vaccine strain, while the addition of the Q189E mutation actually increased virus neutralization by serum antibodies (Figure 7). These data indicate that the E224A and 5a.2a.1 mutations, while potentially deleterious to the virus from an *in vitro* fitness perspective, may have been selected due to their contribution to antigenic drift and evasion of existing immunity. Conversely, the Q189E mutation increases neutralization by serum compared to the vaccine strain whereas the same mutation improved virus fitness *in vitro*. Taken together, these data indicate that the presence of mutations at multiple antigenic sites allows for the tolerance of a mutation, Q189E, that increases serum neutralization, particularly when that mutation can improve *in vitro* virus fitness.

**Figure 7:**
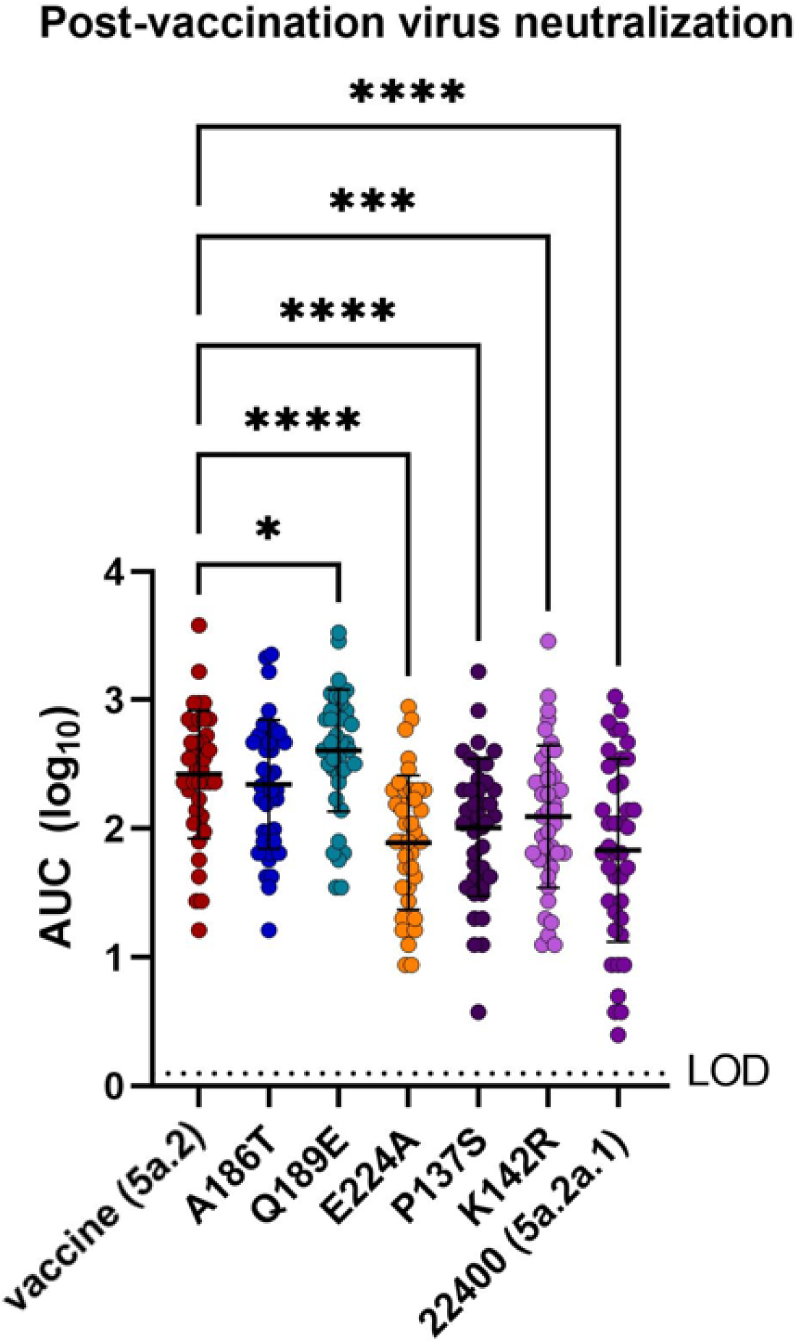
The impact of individual mutations on antigenic drift was assessed on a subset of post-vaccination serum from the 2022-23 Northern Hemisphere influenza vaccine healthcare worker cohort. The E224A, P137S, and K142R mutations reduced virus neutralization by serum antibodies, while the Q189E mutation increased neutralization and A186T had no discernable effect (repeated measures one-way ANOVA with Dunnett post-hoc using A/Victoria as comparison group). N=40. * p<0.05, *** p<0.001, **** p<0.0001

## Discussion

The emergence of the 5a.2a clade of H1N1, and the speed with which it predominated, was surprising considering the failure of its parental clade, 5a.2 (Figure 1A). The 5a.2 clade had significantly reduced *in vitro* fitness compared with the cocirculating 5a.1 clade and their parental 5a clade, including reduced receptor binding diversity, replication, and plaque formation (8). The contrasting success of its progeny 5a.2a clade suggested that there were phenotypic advantages conferred by the 5a.2a clade-defining mutations. Indeed, *in vitro* fitness assays reveal that 5a.2a forms significantly larger plaques (Figure 2A, 2B), replicates significantly better in human nasal epithelial cell (hNEC) cultures (Figure 2C-2F), and has increased receptor binding diversity (Figure 6C) compared with 5a.2. The 5a.2a clade was also antigenically drifted from 5a.2, as demonstrated by neutralization assays using serum from healthcare workers (Figure 3A, 3B). These advantages likely contributed to the ability of 5a.2a to overtake the other co-circulating H1N1 clades.

Three clade-defining mutations for 5a.2a, A186T, Q189E, and E224A, occupy both the receptor binding site (RBS) and antigenic site Sb (Figure 1B), suggesting a key role in fitness differences. When characterized, it was shown that only Q189E was able to independently increase *in vitro* fitness, while E224A actually reduced plaque size and total virus production in hNEC cultures (Figure 5A-F). Interestingly, E224A was the only 5a.2a mutation that contributed to antigenic drift, while Q189E actually resulted in increased virus neutralization (Figure 7). Q189E also appeared as a clade-defining mutation of the 5a.1 clade that predominated from 2020-2021 (Figure 1A) and was present in the 2021-22 Northern Hemisphere (NH) influenza vaccine (46). Exposure to 5a.1 through either natural infection or vaccination may have generated antibody responses against this epitope, and the healthcare workers whose serum was tested above (Figures 3 and 7) would have received the 2021-22 NH vaccine due to workplace vaccination requirements. The 224 residue, on the other hand, has been glutamic acid since the emergence of 2009pdm H1N1 (47), and its switch to alanine in 2021 would have been the first population-level exposure to a new amino acid at this site. The phenotypic contributions of Q189E and E224A highlight the evolutionary balance between mutations that alter fitness and those that contribute to antigenic drift. Given the overlap between the RBS and key antigenic sites, a benefit to one component may come at a cost to the other. The accrual of multiple mutations with contrasting advantages and disadvantages may allow the virus to improve its phenotype while mitigating the deleterious effects of any one change.

Receptor binding specificity is a major factor by which hemagglutinin contributes to virus replication and it has been shown to correlate with viral fitness (5–8). The reduction of *in vitro* fitness of the 5a.2 clade was associated with a loss of receptor binding diversity (8), suggesting that differences in cell attachment may have contributed to the poor circulation of 5a.2. It was therefore hypothesized that the improved *in vitro* fitness of the 5a.2a clade would be associated with an increase in receptor binding diversity, and this was confirmed by glycan microarray experiments (Figure 6). Since the Q189E mutation was shown to be the driver of the improved *in vitro* fitness phenotype, this was the mutation that was expected to contribute to the increased receptor binding diversity. Surprisingly, the effect of Q189E on receptor binding was negligible, while the E224A mutation appears to be the primary cause of the increased diversity (Figure 6). Since the stark effect of Q189E on viral replication and plaque formation was not correlated with an increase in receptor binding diversity, its mechanism for producing the observed *in vitro* fitness changes is not yet completely understood. The glycan microarray used to measure binding diversity consists of synthetic glycans; the mutations tested above may produce differences in receptor binding that are not able to be captured using a synthetic microarray. An array using naturally-derived glycans may reveal differences more specific to the human respiratory tract (48, 49). Glycan microarrays may also overlook differences in low-affinity and non-canonical receptor interactions which have been shown to contribute to viral entry (50, 51). It is also possible that receptor affinity might be altered by one or more of these mutations without producing a significant broadening of receptor binding diversity. Changes in affinity may also explain the differences in replication seen pre-peak versus post-peak for these viruses.

The 224 residue, and specifically the E224A mutation, has previously been studied for its effect on RBS confirmation. It was noted shortly after its emergence that the 2009 pandemic H1N1 may have reduced stability in its interactions between the 190-helix and 220-loop compared with the 1918 H1N1 due to the amino acids at positions 216 and 224 (52). Shang *et al.* modeled differences between these viruses and found that the E224A mutation, in combination with S183P, stabilized interactions between these two regions (27). The addition of E224A to the pdm2009 genetic background was shown to increase binding affinity to fetuin (53) and alter cell tropism (26), though it replicated to lower titers on human airway epithelial cells and had no impact on transmission in ferrets (41). Jayaraman *et al* predicted in 2011 that changing key RBS residues, including E224, may improve receptor binding affinity by creating a more hydrophobic interaction network in this region of the RBS (25, 53, 54). They generated a pdm2009 virus with S183P, A186T, and E224A and demonstrated that it had improved receptor affinity compared with the original pdm2009. Interestingly, all three of these mutations have since become the predominant residues at their respective sites, indicating that this constellation of amino acids is advantageous to the virus. E224A only emerged once these other mutations were also present in circulating H1, and it is possible that there are fitness advantages of E224A that only manifest in the context of these other mutations.

Similarly, while A186T did not appear to produce any apparent *in vitro* fitness advantages on its own, it may produce a synergistic effect when combined with E224A. It has been shown that some mutations only contribute to phenotypic changes when combined with a second mutation, and that the receptor binding phenotype of E224A can be enhanced by additional mutations (25, 53). Experiments that test combinations of recent clade-defining mutations are needed to elucidate any synergistic effects of these amino acid changes.

In addition to assessing differences between the 5a.2a and 5a.2 clades, this study also investigated characteristics of the 5a.2a.1 clade that emerged from 5a.2a. Unlike 5a.2a, 5a.2a.1 did not rapidly predominate within circulating H1N1, but rather gradually increased in prevalence (Figure 1A). The 5a.2a.1 clade was shown to replicate at least as well as 5a.2, and it produced significantly larger plaques and had increased receptor binding diversity (Figures 2 and 6). Both of its major clade-defining mutations, however, resulted in a reduction in performance on one or more *in vitro* fitness assays (Figures 5 and 6). The benefits of these mutations appear to be largely antigenic, as they both significantly reduced serum neutralization when added to the 5a.2 genetic background (Figure 7), and likely contributed to the overall antigenic drift seen for 5a.2a.1 (Figure 3A, 3B).

While the phenotypes of individual 5a.2a.1 mutations were characterized here by adding them to a 5a.2 genetic background, circulating 5a.2a.1 viruses also contain all of the clade-defining mutations associated with 5a.2a. This likely explains the difference in *in vitro* fitness observed between the clinical 5a.2a.1 virus and its individual clade-defining mutations. The deleterious effects of the 5a.2a.1 mutations are likely ameliorated by advantages conferred by one or more 5a.2a mutations, allowing this virus to have adequate fitness while preserving the antigenic changes resulting from the new mutations. This exemplifies the evolutionary balancing act between mutations conferring fitness benefits and those that allow the virus to evade adaptive immunity. One model of influenza evolution proposes the cyclical emergence of mutations with varying levels of receptor avidity influenced by pressure to escape immunity (55), and earlier surveillance of circulating H1N1 since 2009 has shown that mutations increasing receptor avidity were followed by those that reduced avidity (28). An analogous process may be driving recent H1N1 evolution, with the Q189E mutation of 5a.2a increasing viral fitness sufficiently to allow for subsequent 5a.2a.1 mutations driving antigenic drift.

This process can be understood through epistasis, or the impact of secondary mutations on an individual mutant phenotype. Epistatic processes are thought to be a major component of influenza evolution, due in part to overlapping functional regions of hemagglutinin and the frequency of pleiotropic mutations (56, 57). Epistasis in the RBS specifically has been shown to allow for significant sequence diversity and antigenic escape while maintaining viral fitness (58). The striking difference between individual and combined mutations shown here, as well as the recent temporal clustering of RBS mutations, points to a significant epistatic component to contemporary H1N1 clade-defining mutations. The synergistic effect of these mutations, including their contribution to receptor avidity and affinity, should be investigated to determine how the E224A mutation is impacted by the 190-helix mutations, and whether positive epistasis is observed between Q189E and the 5a.2a.1 mutations.

## Methods

### Ethics statement and human subjects

Serum used in this study was obtained from healthcare workers recruited during the annual Johns Hopkins Hospital employee influenza vaccination campaign in the Fall of 2022 by the Johns Hopkins Centers for Influenza Research and Response (JH-CEIRR). Pre- and post-vaccination (∼128 day) human serum were collected from subjects, who provided written informed consent prior to participation. The JHU School of Medicine Institutional Review Board approved this study, IRB00288258. Virus was isolated for this study from deidentified influenza A virus H1N1 positive samples, collected from patients who provided written informed consent during the 2022-23 influenza season at the Johns Hopkins Hospital, under the JHU School of Medicine Institutional Review Board approved protocol, IRB00091667.

### Cell cultures

Madin-Darby canine kidney (MDCK) cells (provided by Dr. Robert A. Lamb) and MDCK-SIAT cells (provided by Dr. Scott Hensley) were maintained in complete medium (CM) consisting of Dulbecco’s Modified Eagle Medium (DMEM) supplemented with 10% fetal bovine serum, 100 units/ml penicillin/streptomycin (Life Technologies) and 2 mM Glutamax (Gibco) at 37 °C and 5% CO_2_. Human nasal epithelial cells (hNEC) (PromoCell) were cultivated as previously described (5, 8).

### Viral RNA Extraction and Multi-segment Amplification by Reverse-Transcriptase Polymerase Chain Reaction

Viral nucleic acid was extracted from samples as previously described (59) using the Chemagic Viral RNA/DNA Kit following the manufacturer’s instructions (Revvity). Samples were eluted into 60μL of elution buffer and used directly after extraction in PCR. The whole genome was amplified using a single multi-segment 1-step reverse-transcription polymerase chain reaction (RT-PCR) with the MBTuni-12 [5’-ACGCGTGATCAGCAAAAGCAGG] and MBTuni-13 [5’-ACGCGTGATCAGTAGAAACAAGG] primer sets for Influenza A (59, 60) with the SuperScript III One-Step RT-PCR kit as per manufacturers guidelines (Invitrogen).

### Library Preparation and Next-Generation Sequencing and Virus Genome Assembly

After amplification, concentrations were checked using the Qubit 2.0 fluorometer (ThermoFisher Scientific). 2μL of PCR product was prepared for sequencing using the native barcoding genomic DNA kit (EXP-NBD196) and the NEBNext ARTIC Library Prep kit, following the manufacturer’s instructions (New England BioLabs / Oxford Nanopore Technologies). 50ng of the barcoded library was sequenced using the R9.4.1 flow cell on a GridION (Oxford Nanopore Technologies). Resulting fastq files were demultiplexed using the artic_guppyplex tool (Artic version 1.2.2). Nucleotide sequence assembly was performed using the FLU module of the Iterative Refinement Meta-Assembler (IRMA version 1.0.2), using IRMA’s default settings, which include a minimum average quality score of 24 and a site depth of 100 (59, 61).

Clinical virus sequences are accessible through GISAID with the following IDs: EPI_ISL_17617226 for A/Baltimore/R0675/2019, EPI_ISL_17626342 for A/Baltimore/JH-22363/2022, EPI_ISL_17626298 for A/Baltimore/JH-22377/2022, and EPI_ISL_17626340 for A/Baltimore/JH-22400/2022.

### Viral isolation

Viral stock preparation was performed as described previously (8, 62) using nasopharyngeal swabs from influenza H1N1-positive individuals during the 2022–2023 Northern Hemisphere influenza season at JHH.

### Virus Cloning and Plasmids

Virus cloning was performed as previously described (5, 10, 63). Viral RNA was extracted using the QIAamp viral RNA mini extraction kit and converted to cDNA using universal influenza A primers. Individual segments were amplified from cDNA with Phusion High-Fidelity DNA Polymerase (New England BioLabs) using segment-specific primers, which also added restriction enzyme sites to the ends of the segments. For all segments, the pHH21 plasmid was used (64, 65). Both the plasmid and individual segments were digested separately with restriction enzymes (either BsmBI or BsaI, New England BioLabs) and then combined and ligated together with T4 DNA ligase (New England BioLabs). Ligated plasmids were transformed into DH5α bacteria (Thermo Fisher) and grown overnight on LB plates containing carbenicillin to select for colonies that successfully received the plasmid. Individual colonies were isolated and grown, and plasmid was extracted using a Qiagen QIAprep Spin Miniprep Kit. Plasmid sequences were verified using restriction enzyme digestion and whole plasmid sequencing, which was performed by Plasmidsaurus using Oxford Nanopore Technology with custom analysis and annotation. Bacteria expressing the correct plasmid were grown in bulk and high-purity plasmid was extracted using ZymoPURE Plasmid Midiprep kit (Zymo research) and verified by sequencing of the coding region of the influenza virus segment.

For generating plasmids containing specific point mutations, mutagenic primers were designed using the New England Biolabs online software NEBaseChanger (primer sequences are available at doi: XXXX. The Q5 Site-Directed Mutagenesis Kit (New England BioLabs) was then used with the A/Baltimore/R0675/2019 hemagglutinin plasmid to produce plasmids with the desired mutations. Plasmids were transformed, extracted, and confirmed as described above.

### Recombinant Virus Production

Recombinant viruses were generated using the 12 plasmid reverse genetics system described previously (5, 44, 45).

### Tissue culture infectious dose 50 (TCID50)

TCID50 assays were performed on MDCK-SIAT cells as previously described (8, 10, 66). The 50% tissue culture infectious dose was calculated as previously described (64, 67).

### Plaque assay

Plaque assays were performed and analyzed as previously described (8, 10) using MDCK cells grown in complete medium to confluence in 6-well plates. Overlay media was phenol-red free DMEM supplemented with .3% BSA (Sigma), 100U/ml pen/strep (Life Technologies), 2mM Glutamax (Gibco), 5mM HEPES buffer (Gibco) 5μg/ml N-acetyl trypsin (Sigma) and 1% cellulose.

### Viral growth curves

hNEC growth curves were performed at an MOI of 0.01 TCID50 units per cell on fully differentiated hNECs grown at an air–liquid interface as described previously (8, 10, 66). Infectious virus concentration in collected media was determined through TCID50.

### Partial virus purification and labeling with Alexa Fluor 488

For each virus analyzed using glycan arrays, 100 mL of virus stock was grown using the method referenced above, and viruses were partially purified through ultracentrifugation and labeled with Alexa Fluor 488 as previously described (8). Labeled virus was stored at −180 °C, and TCID50 was performed to confirm virus activity and concentration. Titer of labeled viruses were: 4.20E+8 TCID50/mL (A/Baltimore/R0675/2019), 1.33E+8 TCID50/mL (A/Baltimore/JH-22363/2022), 1.70E+8 TCID50/mL (A/Baltimore/JH-22377/2022), 5.37E+8 TCID50/mL (A/Baltimore/JH-22400), 1.58E+8 TCID50/mL (A/Baltimore/R0675/2019 with A186T), 1.95E+8 TCID50/mL (A/Baltimore/R0675/2019 with Q189E), 4.20E+7 TCID50/mL (A/Baltimore/R0675/2019 with E224A), 6.17E+7 TCID50/mL (A/Baltimore/R0675/2019 with P137S), and 2.86E+7 TCID50/mL (A/Baltimore/R0675/2019 with K142R).

### Glycan array

The Consortium for Functional Glycomics version 5.5 microarray, containing 562 glycans in replicates of 6, was used for glycan arrays as previously described (8, 68). Microarray slides were submerged in 100 ml of PBS wash buffer (phosphate-buffered saline containing 0.005% Tween-20) for five minutes to re-hydrate. 70 µl of undiluted sample were loaded onto the slide. A cover slip was placed over the slide and incubated at room temperature for one hour in darkness, after which the cover slip was removed and the slide was washed four times in 100 ml of PBS wash buffer, then PBS, then deionized water. Slides were then dried before being read in a GenePix microarray scanner.

For each set of six replicates, only the median four values were used in analysis. A graph of the raw data was visually analyzed to confirm selective binding of terminally sialylated glycans, after which data was filtered to only include these glycans for a total of 139 glycans. Data was then normalized to a maximum value of 100, and organized by structure using the Glycan Array Dashboard (GLAD) (69). Both the heat map and structures of individual glycans were generated through GLAD.

### Serum neutralization assay

Pre- and post-vaccination human serum was treated with Receptor Destroying Enzyme II (RDE, Hardy Diagnostics) and neutralization assays were performed on MDCK-SIAT cells as previously described (8, 70), performed in quadruplicate and using 100 TCID50 of virus per well.

### Statistical analyses

All statistical analyses were performed in GraphPad Prism 9.1.0. Growth curves were analyzed using 2-way ANOVA with Tukey post-hoc test. Area under the curve (AUC) was calculated for each growth curve replicate and AUC values were compared using one-way ANOVA for differences in total virus production. Plaque assays were analyzed using Kruskal–Wallis ANOVA with Dunn’s multiple comparison test. Neutralization assays and fold-change in neutralizing titers were analyzed using repeated-measures one-way ANOVA with Tukey post-hoc, and differences in seroconversion between clades were analyzed using a Chi-square test.

## Data availability

The datasets used and/or analyzed in the current study are available through the Johns Hopkins Research Data Repository at doi: XXXX

## Acknowledgements

This work was supported by National Institutes of Health (NIH) contracts N272201400007C and N75 N03021C00045 for the Johns Hopkins Centers of Excellence in Influenza Research and Research, NIH T32 AI007417, NIH U54 AG062333, as well as the Richard Eliasberg Family Foundation. We acknowledge the Protein-Glycan Interaction Resource of the CFG and the National Center for Functional Glycomics (NCFG) at Beth Israel Deaconess Medical Center, Harvard Medical School (supporting grant R24GM137763) for glycan microarray studies, as well as NextStrain.org for their open access influenza surveillance dashboard, PyMOL molecular visualization software for use in mapping hemagglutinin mutants, and Plasmidsaurus for plasmid sequencing. The authors thank the healthcare workers who enrolled and participated in the study. We are grateful for the efforts of the clinical coordination team at JHH who collected samples. We thank the laboratories of Jenna Guthmiller, Sabra Klein, Kimberly Davis, Nicole Baumgarth, and Andrew Pekosz for discussion of data and future directions.

## References

1. WHO. 4 January 2023 2018. Influenza (seasonal), on World Health Organization. https://www.who.int/en/news-room/fact-sheets/detail/influenza-(seasonal). Accessed

2. Paul Glezen W, Schmier JK, Kuehn CM, Ryan KJ, Oxford J. 2013. The burden of influenza B: a structured literature review. American journal of public health 103:e43–e51.

3. Tokars JI, Olsen SJ, Reed C. 2018. Seasonal incidence of symptomatic influenza in the United States. Clinical Infectious Diseases 66:1511–1518.

4. Petrova VN, Russell CA. 2018. The evolution of seasonal influenza viruses. Nature Reviews Microbiology 16:47–60.

5. Powell H, Liu H, Pekosz A. 2021. Changes in sialic acid binding associated with egg adaptation decrease live attenuated influenza virus replication in human nasal epithelial cell cultures. Vaccine 39:3225–3235.

6. de Graaf M, Fouchier RA. 2014. Role of receptor binding specificity in influenza A virus transmission and pathogenesis. The EMBO journal 33:823–841.

7. Nobusawa E, Ishihara H, Morishita T, Sato K, Nakajima K. 2000. Change in receptor-binding specificity of recent human influenza A viruses (H3N2): a single amino acid change in hemagglutinin altered its recognition of sialyloligosaccharides. Virology 278:587–596.

8. Swanson NJ, Marinho P, Dziedzic A, Jedlicka A, Liu H, Fenstermacher K, Rothman R, Pekosz A. 2023. 2019–2020 H1N1 clade A5a. 1 viruses have better in vitro fitness compared with the co-circulating A5a. 2 clade. Scientific Reports 13:10223.

9. Wohlgemuth N, Ye Y, Fenstermacher KJ, Liu H, Lane AP, Pekosz A. 2017. The M2 protein of live, attenuated influenza vaccine encodes a mutation that reduces replication in human nasal epithelial cells. Vaccine 35:6691–6699.

10. Powell H, Pekosz A. 2020. Neuraminidase antigenic drift of H3N2 clade 3c. 2a viruses alters virus replication, enzymatic activity and inhibitory antibody binding. PLoS pathogens 16:e1008411.

11. Conenello GM, Tisoncik JR, Rosenzweig E, Varga ZT, Palese P, Katze MG. 2011. A single N66S mutation in the PB1-F2 protein of influenza A virus increases virulence by inhibiting the early interferon response in vivo. Journal of virology 85:652–662.

12. Wei K, Sun H, Sun Z, Sun Y, Kong W, Pu J, Ma G, Yin Y, Yang H, Guo X. 2014. Influenza A virus acquires enhanced pathogenicity and transmissibility after serial passages in swine. Journal of virology 88:11981–11994.

13. Meng F, Yang H, Qu Z, Chen Y, Zhang Y, Zhang Y, Liu L, Zeng X, Li C, Kawaoka Y. 2022. A Eurasian avian-like H1N1 swine influenza reassortant virus became pathogenic and highly transmissible due to mutations in its PA gene. Proceedings of the National Academy of Sciences 119:e2203919119.

14. Hu J, Hu Z, Wei Y, Zhang M, Wang S, Tong Q, Sun H, Pu J, Liu J, Sun Y. 2022. Mutations in PB2 and HA are crucial for the increased virulence and transmissibility of H1N1 swine influenza virus in mammalian models. Veterinary Microbiology 265:109314.

15. Griffin EF, Tompkins SM. 2023. Fitness Determinants of Influenza A Viruses. Viruses 15:1959.

16. Merced-Morales A, Daly P, Abd Elal AI, Ajayi N, Annan E, Budd A, Barnes J, Colon A, Cummings CN, Iuliano AD. 2022. Influenza activity and composition of the 2022–23 influenza vaccine— United States, 2021–22 season. Morbidity and Mortality Weekly Report 71:913.

17. Skehel JJ, Wiley DC. 2000. Receptor binding and membrane fusion in virus entry: the influenza hemagglutinin. Annual review of biochemistry 69:531–569.

18. WHO. 24 September 2021 2021. Recommended composition of influenza virus vaccines for use in the 2022 southern hemisphere influenza season. https://www.who.int/publications/m/item/recommended-composition-of-influenza-virus-vaccines-for-use-in-the-2022-southern-hemisphere-influenza-season. Accessed

19. Zolotarova O, Budzanivska I, Leibenko L, Radchenko L, Mironenko A. 2019. Antigenic site variation in the hemagglutinin of pandemic influenza A (H1N1) pdm09 viruses between 2009– 2017 in Ukraine. Pathogens 8:194.

20. El Moussi A, Ben Hadj Kacem MA, Pozo F, Ledesma J, Cuevas MT, Casas I, Slim A. 2013. Genetic diversity of HA1 domain of heammaglutinin gene of influenza A (H1N1) pdm09 in Tunisia. Virology journal 10:1–9.

21. Guldemir D, Kalaycioglu AT, Altas AB, Korukluoglu G, Durmaz R. 2013. Monitoring genetic diversity of influenza A (H1N1) pdm09 virus circulating during the post-pandemic period in Turkey. Japanese journal of infectious diseases 66:299–305.

22. Ledesma J, Pozo F, Reina G, Blasco M, Rodríguez G, Montes M, López-Miragaya I, Salvador C, Reina J, de Lejarazu RO. 2012. Genetic diversity of influenza A (H1N1) 2009 virus circulating during the season 2010–2011 in Spain. Journal of clinical virology 53:16–21.

23. Ramos AP, Herrera BA, Ramírez OV, García AA, Jiménez MM, Valdés CS, Fernández AG, González G, Fernández SIO, Báez GG. 2013. Molecular and phylogenetic analysis of influenza A H1N1 pandemic viruses in Cuba, May 2009 to August 2010. International Journal of Infectious Diseases 17:e565–e567.

24. Wedde M, Biere B, Wolff T, Schweiger B. 2015. Evolution of the hemagglutinin expressed by human influenza A (H1N1) pdm09 and A (H3N2) viruses circulating between 2008–2009 and 2013–2014 in Germany. International Journal of Medical Microbiology 305:762–775.

25. Martínez-Romero C, de Vries E, Belicha-Villanueva A, Mena I, Tscherne DM, Gillespie VL, Albrecht RA, de Haan CA, García-Sastre A. 2013. Substitutions T200A and E227A in the hemagglutinin of pandemic 2009 influenza A virus increase lethality but decrease transmission. Journal of virology 87:6507–6511.

26. van Doremalen N, Shelton H, Roberts KL, Jones IM, Pickles RJ, Thompson CI, Barclay WS. 2011. A single amino acid in the HA of pH1N1 2009 influenza virus affects cell tropism in human airway epithelium, but not transmission in ferrets. PLoS One 6:e25755.

27. Shang C, Whittleston CS, Sutherland-Cash KH, Wales DJ. 2015. Analysis of the Contrasting Pathogenicities Induced by the D222G Mutation in 1918 and 2009 Pandemic Influenza A Viruses. Journal of Chemical Theory and Computation 11:2307–2314.

28. de Vries RP, de Vries E, Martínez-Romero C, McBride R, van Kuppeveld FJ, Rottier PJ, García-Sastre A, Paulson JC, de Haan CA. 2013. Evolution of the hemagglutinin protein of the new pandemic H1N1 influenza virus: maintaining optimal receptor binding by compensatory substitutions. Journal of virology 87:13868–13877.

29. Martínez-Baz I, Fernández-Huerta M, Navascués A, Pozo F, Trobajo-Sanmartín C, Casado I, Echeverria A, Ezpeleta C, Castilla J. 2023. Influenza vaccine effectiveness in preventing laboratory-confirmed influenza cases and hospitalizations in Navarre, Spain, 2022–2023. Vaccines 11:1478.

30. WHO. 23 September 2022 2022. Recommended composition of influenza virus vaccines for use in the 2023 southern hemisphere influenza season. https://www.who.int/publications/m/item/recommended-composition-of-influenza-virus-vaccines-for-use-in-the-2023-southern-hemisphere-influenza-season. Accessed

31. Opanda S, Bulimo W, Gachara G, Ekuttan C, Amukoye E. 2020. Assessing antigenic drift and phylogeny of influenza A (H1N1) pdm09 virus in Kenya using HA1 sub-unit of the hemagglutinin gene. PLoS One 15:e0228029.

32. Monamele CG, Munshili Njifon HL, Vernet M-A, Njankouo MR, Kenmoe S, Yahaya AA, Deweerdt L, Nono R, Mbacham W, Anong DN. 2019. Molecular characterization of influenza A (H1N1) pdm09 in Cameroon during the 2014-2016 influenza seasons. PloS one 14:e0210119.

33. Kao C-L, Chan T-C, Tsai C-H, Chu K-Y, Chuang S-F, Lee C-C, Li Z-RT, Wu K-W, Chang L-Y, Shen Y-H. 2012. Emerged HA and NA mutants of the pandemic influenza H1N1 viruses with increasing epidemiological significance in Taipei and Kaohsiung, Taiwan, 2009–10. PLoS One 7:e31162.

34. Tapia R, Torremorell M, Culhane M, Medina RA, Neira V. 2020. Antigenic characterization of novel H1 influenza A viruses in swine. Scientific reports 10:4510.

35. Ortiz L, Geiger G, Ferreri L, Moran D, Alvarez D, Gonzalez-Reiche AS, Mendez D, Rajao D, Cordon-Rosales C, Perez DR. 2023. Evolution and Introductions of Influenza A Virus H1N1 in a Farrow-to-Finish Farm in Guatemala. Microbiology Spectrum 11:e02878–22.

36. Kowalczyk A, Urbaniak K, Markowska-Daniel I, Pejsak Z. 2015. Substitution in position 222 of haemagglutinin of pandemic influenza A (H1N1) viruses isolated from pigs in Poland. Journal of Veterinary Research 59:451–456.

37. Markin A, Ciacci Zanella G, Arendsee ZW, Zhang J, Krueger KM, Gauger PC, Vincent Baker AL, Anderson TK. 2023. Reverse-zoonoses of 2009 H1N1 pandemic influenza A viruses and evolution in United States swine results in viruses with zoonotic potential. PLoS Pathogens 19:e1011476.

38. Ibricevic A, Pekosz A, Walter MJ, Newby C, Battaile JT, Brown EG, Holtzman MJ, Brody SL. 2006. Influenza virus receptor specificity and cell tropism in mouse and human airway epithelial cells. Journal of virology 80:7469–7480.

39. Fischer W. 2015. II; King, LS; Lane, AP; Pekosz, A. Restricted replication of the live attenuated influenza a virus vaccine during infection of primary differentiated human nasal epithelial cells. Vaccine 33:4495–4504.

40. Forero A, Fenstermacher K, Wohlgemuth N, Nishida A, Carter V, Smith EA, Peng X, Hayes M, Francis D, Treanor J. 2017. Evaluation of the innate immune responses to influenza and live-attenuated influenza vaccine infection in primary differentiated human nasal epithelial cells. Vaccine 35:6112–6121.

41. Liu H, Gong Y-N, Shaw-Saliba K, Mehoke T, Evans J, Liu Z-Y, Lewis M, Sauer L, Thielen P, Rothman R. 2021. Differential disease severity and whole-genome sequence analysis for human influenza A/H1N1pdm virus in 2015–2016 influenza season. Virus evolution 7:veab044.

42. van Riel D, den Bakker MA, Leijten LM, Chutinimitkul S, Munster VJ, de Wit E, Rimmelzwaan GF, Fouchier RA, Osterhaus AD, Kuiken T. 2010. Seasonal and pandemic human influenza viruses attach better to human upper respiratory tract epithelium than avian influenza viruses. The American journal of pathology 176:1614–1618.

43. Lee C-W. 2014. Reverse genetics of influenza virus. Animal influenza virus:37–50.

44. Neumann G, Watanabe T, Ito H, Watanabe S, Goto H, Gao P, Hughes M, Perez DR, Donis R, Hoffmann E. 1999. Generation of influenza A viruses entirely from cloned cDNAs. Proceedings of the National Academy of Sciences 96:9345–9350.

45. Fodor E, Devenish L, Engelhardt OG, Palese P, Brownlee GG, García-Sastre A. 1999. Rescue of influenza A virus from recombinant DNA. J Virol 73:9679–82.

46. WHO. 28 February 2020 2020. Recommended composition of influenza virus vaccines for use in the 2020-2021 northern hemisphere influenza season. https://www.who.int/publications/m/item/recommended-composition-of-influenza-virus-vaccines-for-use-in-the-2020-2021-northern-hemisphere-influenza-season. Accessed

47. Hadfield J, Megill C, Bell SM, Huddleston J, Potter B, Callender C, Sagulenko P, Bedford T, Neher RA. 2018. Nextstrain: real-time tracking of pathogen evolution. Bioinformatics 34:4121–4123.

48. Byrd-Leotis L, Gao C, Jia N, Mehta AY, Trost J, Cummings SF, Heimburg-Molinaro J, Cummings RD, Steinhauer DA. 2019. Antigenic pressure on H3N2 influenza virus drift strains imposes constraints on binding to sialylated receptors but not phosphorylated glycans. Journal of virology 93:e01178–19.

49. McQuillan AM, Byrd-Leotis L, Heimburg-Molinaro J, Cummings RD. 2019. Natural and synthetic sialylated glycan microarrays and their applications. Frontiers in Molecular Biosciences 6:88.

50. Glaser L, Conenello G, Paulson J, Palese P. 2007. Effective replication of human influenza viruses in mice lacking a major α2, 6 sialyltransferase. Virus research 126:9–18.

51. Liu M, Huang LZ, Smits AA, Büll C, Narimatsu Y, van Kuppeveld FJ, Clausen H, de Haan CA, de Vries E. 2022. Human-type sialic acid receptors contribute to avian influenza A virus binding and entry by hetero-multivalent interactions. Nature Communications 13:1–12.

52. Maines TR, Jayaraman A, Belser JA, Wadford DA, Pappas C, Zeng H, Gustin KM, Pearce MB, Viswanathan K, Shriver ZH. 2009. Transmission and pathogenesis of swine-origin 2009 A (H1N1) influenza viruses in ferrets and mice. Science 325:484–487.

53. de Vries RP, de Vries E, Moore KS, Rigter A, Rottier PJ, de Haan CA. 2011. Only two residues are responsible for the dramatic difference in receptor binding between swine and new pandemic H1 hemagglutinin. Journal of Biological Chemistry 286:5868–5875.

54. Jayaraman A, Pappas C, Raman R, Belser JA, Viswanathan K, Shriver Z, Tumpey TM, Sasisekharan R. 2011. A single base-pair change in 2009 H1N1 hemagglutinin increases human receptor affinity and leads to efficient airborne viral transmission in ferrets. PloS one 6:e17616.

55. Hensley SE, Das SR, Bailey AL, Schmidt LM, Hickman HD, Jayaraman A, Viswanathan K, Raman R, Sasisekharan R, Bennink JR. 2009. Hemagglutinin receptor binding avidity drives influenza A virus antigenic drift. Science 326:734–736.

56. Lyons DM, Lauring AS. 2018. Mutation and epistasis in influenza virus evolution. Viruses 10:407.

57. Kryazhimskiy S, Dushoff J, Bazykin GA, Plotkin JB. 2011. Prevalence of epistasis in the evolution of influenza A surface proteins. PLoS genetics 7:e1001301.

58. Wu NC, Xie J, Zheng T, Nycholat CM, Grande G, Paulson JC, Lerner RA, Wilson IA. 2017. Diversity of functionally permissive sequences in the receptor-binding site of influenza hemagglutinin. Cell host & microbe 21:742–753. e8.

59. Fall A, Han L, Yunker M, Gong Y-N, Li T-J, Norton JM, Abdullah O, Rothman RE, Fenstermacher KZ, Morris CP. Evolution of influenza A (H3N2) viruses in 2 consecutive seasons of genomic surveillance, 2021–2023, pofad577. In (ed), Oxford University Press US,

60. Zhou B, Donnelly ME, Scholes DT, St. George K, Hatta M, Kawaoka Y, Wentworth DE. 2009. Single-reaction genomic amplification accelerates sequencing and vaccine production for classical and Swine origin human influenza a viruses. Journal of virology 83:10309–10313.

61. Fall A, Gallagher N, Morris CP, Norton JM, Pekosz A, Klein E, Mostafa HH. 2022. Circulation of enterovirus D68 during period of increased influenza-like illness, Maryland, USA, 2021. Emerging infectious diseases 28:1525.

62. Blumenkrantz DR, Mehoke T, Shaw-Saliba K, Powell H, Wohlgemuth N, Liu H, Macias E, Evans J, Lewis M, Medina R. 2021. Identification of H3N2 NA and PB1-F2 genetic variants and their association with disease symptoms during the 2014–15 influenza season. Virus Evolution 7:veab047.

63. Klein EY, Blumenkrantz D, Serohijos A, Shakhnovich E, Choi J-M, Rodrigues JV, Smith BD, Lane AP, Feldman A, Pekosz A. 2018. Stability of the influenza virus hemagglutinin protein correlates with evolutionary dynamics. Msphere 3:10.1128/mspheredirect.00554-17.

64. McCown MF, Pekosz A. 2005. The influenza A virus M2 cytoplasmic tail is required for infectious virus production and efficient genome packaging. Journal of virology 79:3595–3605.

65. McCown MF, Pekosz A. 2006. Distinct domains of the influenza a virus M2 protein cytoplasmic tail mediate binding to the M1 protein and facilitate infectious virus production. Journal of virology 80:8178–8189.

66. Resnick JD, Beer MA, Pekosz A. 2023. Early transcriptional responses of human nasal epithelial cells to infection with Influenza A and SARS-CoV-2 virus differ and are influenced by physiological temperature. Pathogens 12:480.

67. Reed LJ, Muench H. 1938. A simple method of estimating fifty per cent endpoints. American journal of epidemiology 27:493–497.

68. Heimburg-Molinaro J, Tappert M, Song X, Lasanajak Y, Air G, Smith DF, Cummings RD. 2012. Probing virus–glycan interactions using glycan microarrays, p 251–267, Carbohydrate Microarrays. Springer.

69. Mehta AY, Cummings RD. 2019. GLAD: GLycan Array Dashboard, a visual analytics tool for glycan microarrays. Bioinformatics 35:3536–3537.

70. Wilson JL, Akin E, Zhou R, Jedlicka A, Dziedzic A, Liu H, Fenstermacher KZ, Rothman RE, Pekosz A. 2023. The Influenza B Virus Victoria and Yamagata Lineages Display Distinct Cell Tropism and Infection-Induced Host Gene Expression in Human Nasal Epithelial Cell Cultures. Viruses 15:1956.

